# Immunoprofiling of HTLV-1-infected individuals shows altered innate cell responsiveness in HAM/TSP patients

**DOI:** 10.1101/2021.04.07.438775

**Authors:** Brenda Rocamonde, Nicolas Futsch, Noemia Orii, Omran Allatif, Augusto Cesar Penalva de Oliveira, Renaud Mahieux, Jorge Caseb, Hélène Dutartre

**Affiliations:** International Center for Research in Infectiology, Retroviral Oncogenesis Laboratory, INSERM U1111 - Université Claude Bernard Lyon 1, CNRS, UMR5308, Ecole Normale Supérieure de Lyon, Université Lyon, Lyon, France, Equipe labelisée par la Fondation pour la Recherche Médicale, Labex Ecofect; Faculdade de Medicina/ Instituto de Medicina Tropical de São Paulo/Universidade da São Paulo, São Paulo, SP, Brazil; International Center for Research in Infectiology, service BIBS, INSERM U1111 - Université Claude Bernard Lyon 1, CNRS, UMR5308, Ecole Normale Supérieure de Lyon, Université Lyon, Lyon, France; Instituto de Infectologia “Emilio Ribas”, São Paulo, SP, Brazil

## Abstract

The Human T-cell Leukemia Virus-1 (HTLV-1)-Associated Myelopathy/Tropical Spastic Paraparesis (HAM/TSP) is a devastating neurodegenerative disease with no effective treatment, which affects an increasing number of people in Brazil. A biological blood factor allowing the prediction of the disease occurrence is so far not available. In this study, we analyzed innate immunity responses at steady state and after blood cell stimulation using an agonist of the toll-like receptor (TLR)7/8-signaling pathway in blood samples from HTLV-1-infected volunteers, including asymptomatic carriers and HAM/TSP patients. We observed a lower responsiveness in dendritic cells to produce IFNα. Moreover, we found higher production of IL-12 and Mip-1α by monocytes together with higher levels of IFNγ produced by Natural Killer cells. These deregulations could represent a signature for progression towards HAM/TSP.

## Introduction

The Human T-cell Leukemia Virus-1 (HTLV-1)-Associated Myelopathy/Tropical Spastic Paraparesis (HAM/TSP)^1,2^ is a progressive neurodegenerative disease characterized by the demyelination of the middle-to-lower thoracic cord^1^, this illness presenting a high prevalence in Brazil^3^. Even though most HTLV-1 infected individuals remain asymptomatic lifelong, around 1-5% develop HAM/TSP. However, anticipating those infected asymptomatic carriers who will develop HAM/TSP remains a challenging task. Viral and immune characterization of HAM/TSP patients identify some markers of pathogenicity, such as an increased proviral load (PVL, *i*.*e*. the number of integrated copies of the viral genome in the host cells) or higher frequency of CD8^+^ T-cells and higher secretion of pro-inflammatory cytokines such as TNFα and IFNγ^4^. However, those markers do not allow to anticipate the development of HAM/TSP in infected asymptomatic carriers, since some of them also present an elevated PVL without developing any myelopathy, and alterations in T-cell frequencies and cytokine secretion is only detectable in patients with disease manifestations, and not in infected asymptomatic carriers^5^. Thus, new markers of the disease progression remain to be identified.

HTLV-1 targets mainly T-cells^6^, altering their function and ability to induce an antiviral specific immune response, even participating in the disease evolution as mentioned previously. However, in addition to T-cells, HTLV-1 also targets innate immune cells such as classical and plasmacytoid dendritic cells (cDCs and pDCs, respectively)^7–9^ as well as monocytes^10^. Yet, their role in the disease manifestation is poorly understood, besides alterations in innate cell frequencies^8,10–12^ observed in HAM/TSP patients. Indeed, the frequencies of pDCs^13^ were found lower while those of myeloid DCs^13^ and of intermediate monocytes^12^ were found higher in HAM/TSP patients compared to HTLV-1 asymptomatic carriers and healthy donors. In contrast, while higher frequencies of non-classical monocytes and lower frequencies of classical monocyte were reported in HTLV-1-infected individuals compared to healthy donors^10^, these frequencies were not different between infected asymptomatic carriers and HAM/TSP patients^10^. Strikingly, the innate cells responsiveness has not been addressed yet, although a dysfunctional immune response linked either to the infection of innate cells or to their activation upon virus sensing, might be an underlying mechanism involved in HAM/TSP progression. Therefore, innate immune responsiveness and its potential deregulation after HTLV-1 infection require special attention to understand HAM/TSP pathology and disease progression.

The aim of this study was to investigate potential deregulations in innate cell responsiveness in HTLV-1 infected subjects that could indicate a progression towards HAM/TSP. We performed single cell immunoprofiling of freshly collected blood samples from a cohort of asymptomatic carriers and HAM/TSP patients to characterize the phenotype and responsiveness of innate cell subsets after *ex vivo* TLR7/8-stimulation, a broad way to activate most of innate cells^14^ and because, TLR7/8 signaling impairment upon several chronic infections has been linked to diseases^15,16^. Up-to-now, the increase in PVL is the only factor to predict progression of asymptomatic infection to disease, but this cannot anticipate whether chronic HTLV-1-infection will evolve to HAM/TSP pathology. Thus we have evaluated the alterations in cell frequency but also the immune response of a cohort of 30 HTLV-1-infected Brazilians –considering both HAM/TSP and asymptomatic carriers– and 15 age and sex-paired individuals as controls. We have reported the presence of TNFα-producing clusters in asymptomatic carriers, diminished in HAM/TSP donors possibly due to corticosteroid treatment at the moment of the analysis. However, TLR7/8 stimulation evinced a pro-inflammatory signature in HAM/TSP monocytes with higher production of IL-12 and Mip-1α. Moreover, the IFNα response driven by dendritic cells was significantly lower in HAM/TSP patients, suggesting an impaired antiviral response to the infection.

### Methodological approach

#### Clinical samples

A Brazilian cohort (15 HAM/TSP patients, 15 HTLV-1 asymptomatic carriers, and 15 non-infected individuals) was analyzed. All individuals were followed at Institute of Infectious Diseases “Emílio Ribas” (IIER) and signed an informed consent that was approved by the local Ethical Board at the Institute of Infectious Diseases “Emílio Ribas”. Patients underwent a neurological assessment by a neurologist blinded to their HTLV status. Patients with at least two pyramidal signs, such as paresis, spasticity, hyperreflexia, clonus, diminished or absent superficial reflexes, or the presence of pathologic reflexes (e.g. Babinski sign), as defined in Castro-Costa Criteria, 2006, were defined as having HAM/TSP and all diagnosed HAM/TSP received corticosteroid treatment (prednisolone) 45 days apart. Asymptomatic carriers were included based on their HTLV-1 positive status and their lack of any HTLV-1 associated clinical symptoms. They were aged and sex matched with the HAM/TSP patients enrolled. Detailed clinical information from HTLV-1 infected individuals included in the cohort is provided in Supplementary Table 1.

### HTLV-1 serologic test and proviral load (PVL) determination

HTLV-1 serologic diagnosis was made by ELISA (Ortho Diagnostics, USA) and positive samples were confirmed by western blot (HTLV Blot 2.4 test, DBL, Singapore). All patients whose serum sample was reactive with either test was submitted to a nested-PCR using HTLV-1 generic primers and amplified products were digested with restriction enzymes^17^. In order to determine HTLV-1 proviral load, peripheral blood mononuclear cells (PBMC) were isolated from an acid-citrate-dextrose solution and separated by Ficoll density gradient centrifugation (Pharmacia, Uppsala, Sweden). DNA was extracted using a commercial kit (GFX Pharmacia, Uppsala, Sweden). The forward and reverse primers used for HTLV-1 DNA quantitation were SK110 (5’-CCCTACAATCCAACCAGCTCAG-3’, HTLV-1 nucleotide 4758-4779 (GenBank accession No. J02029)), and SK111 (5’-GTGGTGAAGCTGCCATCGGGTTTT-3’, HTLV-1 nucleotide 4943-4920). The internal HTLV-1 TaqMan probe (5’-CTTTAC TGACAAACCCGACCTACCCATGGA-3’) was selected using the Oligo (version 4, National Biosciences, Plymouth, MI, USA) and Primer Express (Perkin-Elmer Applied Biosystems, Boston, MA, USA) software programs. The probe was located between positions 4829 and 4858 of the HTLV-1 genome and carried a 5’ reporter dye FAM (6-carboxy fluorescein) and a 3’ quencher dye TAMRA (6-carboxy tetramethyl rhodamine). Albumin DNA quantification was used to normalize variations due to differences of DNA extraction or PBMCs counts as described previously ^18^. The normalized value of HTLV-1 proviral load was calculated as the ratio of (HTLV-1 DNA average copy number/albumin DNA average copy number) x 2 × 10^6^ and is reported as the number of HTLV-1 copies/10^5^ PBMC^19^.

### Whole blood stimulation

Collected blood samples were distributed in 1.5 mL polypropylene tubes and supplemented with 200 µL of RPMI medium containing 10% of fetal bovine serum (FBS). Samples were cultured in the presence of Resiquimod (R848, 1 µg/mL, Invivogen) to simulate TLR7 signaling pathway. Samples cultured in absence of stimulus were used as controls. After 1h of incubation at 37°C 5% CO_2_, Brefeldine A (10 µg/mL, Sigma) was added to repress cytokine release. Four hours later, samples were incubated for 10 minutes with ammonium chloride in order to perform the lysis of red cells. Staining was performed after one wash with DPBS 1x (Gibco) on whole leukocytes.

### Phenotypic characterization

Samples were incubated with a Live Dead Aqua Blue reagent (Thermo Fisher Scientific) according to manufacturer instructions. After one wash, cells were saturated with 1% BSA-FcR Blocking (Miltenyi) in DPBS for 15 min at 4°C, and then surface-stained for 20 min at 4°C with a cocktail of coupled-antibodies (Supplementary Table 2A). Leukocytes were then fixed for 20 min at room temperature with 4% paraformaldehyde, permeabilized with 0.05% Saponine-DPBS, and stained with coupled-antibodies directed against intracellular cytokines (Supplementary Table 2B). Samples were finally analyzed with a LSR Fortessa X-20 cytometer (BD Bioscience). Fluorochrome compensation was performed with compensation beads (BD Bioscience) and FMO (Fluorescence Minus One) conditions.

### Gating strategy

All data were analyzed using FlowJo™ Software Version 10.5.3 for Mac OS X. Major lineage subsets were selected from forward and side scatter properties followed by single live cells (Supplementary Figure 1). Doublet discrimination was achieved by plotting FSC-H vs. SSC-A. For innate immunity live, single, HLA-DR+ cells were selected. Hierarchical gating allows then the discrimination of the following innate cell subsets: cDC1 (CD11c^+^, BDCA2/3^+^), cDC2 (CD11c^+^, BDCA2/3^-^, BDCA1^+^), pDC (CD11c^-^, BDCA2/3^+^), monocytes (CD11c+, BDCA2/3^-^, BDCA1^-^). CD16 and CD14 expression further defined the following subsets of monocytes: classical (CD14^+^CD16^-^), intermediate (CD14^+^CD16^+^) and non-classical monocytes (CD14^-^CD16^+^). NK populations were defined as HLA-DR^-^, Lin^-^, and subdivided into CD56^dim^CD16^+^ and CD56^high^CD16^-^ NK cell subsets. For the analysis of cytokine production, positive populations were determined after gating determined using fluorescence Minus One (FMO), and the complete gating from innate cell responsiveness of one representative sample is shown in Supplemental Figure 2. Gating was applied to all samples and was manually checked for consistency across all samples.

### Biostatistical and computational analyses

#### (i) Biostatistical analysis

Biostatistical analysis and data processing were performed using the R programming language. In order to determine statistically significant differences between clinical groups, a test on the homogeneity of variances across samples was applied first (Bartlett’s test). One-way ANOVA was performed when H0 (= Equal variances) was not rejected, followed by Turkey post-hoc test. Otherwise non-parametric ANOVA (*i*.*e*. Kruskal-Wallis test) was applied. To note, logarithmic transformation of factors not following a normal distribution did not improve statistical performances.

#### (ii) tSNE analysis

T-distributed stochastic neighborhood embedding (tSNE) analysis was performed with FlowJo™ Software Version 10.5.3 for Mac OS X. cDC11^+^ and BDCA2-3^+^ cells were selected for the analysis and a downsampling of 3000 cells was performed to have the same number of cells per subject. Only surface markers were selected to perform tSNE analysis with 1000 iterations and a perplexity of 20.

## Results

### Sociodemographic factors or PVL do not predict a HAM/TSP progression

A total of 45 age-matched individuals were enrolled in the study. In order to match the reported higher prevalence of HTLV-1 infection among women^20^, we included twice/three times more women than men. The age, sex, PVL, the clinical motor score and some other information about risk factors such as prolonged breastfeeding is detailed in Supplementary Table 1. The average age was 52 years (±6.35) for men and 49.15 years (±10.84) for women. Age means were 49.8 years for healthy donors (HD), 46.93 years for asymptomatic carriers (AC) and 53.8 years for HAM/TSP. Although an increased PVL was considered the only hallmark of HAM/TSP^21^, we found no significance differences between AC and HAM/TSP subjects (Figure 1A).

We then wondered whether there was a correlation between the proviral charge and motor dysfunction indicators. However, we found no direct correlation between the PVL and either IPEC or Osame score of the HAM/TSP patients (Figure 1B), suggesting that PVL is not a fair indicator of the disability/disease degree. Interestingly, a correlation was found between the PVL and the age of the patients (Figure 1C). Suggesting that increased PVL would be a result of accumulative viral exposition or aged-related factors rather than an indicator of the disease onset.

**Figure 1.**
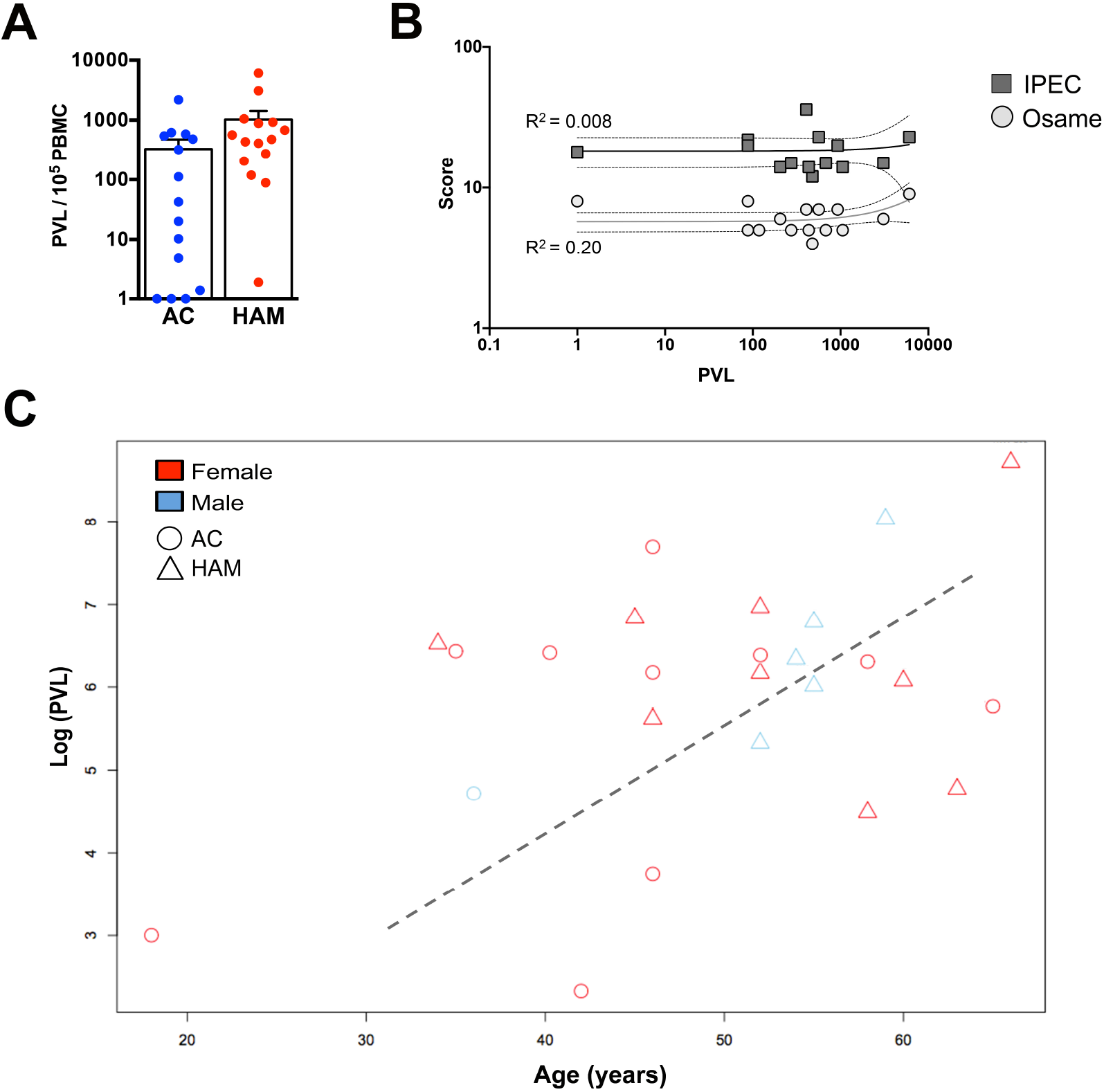
**(A)** Proviral load (PVL) of the HTLV-1 asymptomatic carriers (AC) and HAM/TSP patients. **(B)** Motor score (IPEC and Osame) do not correlate with higher PVL in HAM/TSP patients. **(C)** PVL of HTLV-1-infected individuals shows a correlation between the PVL and the age.

### Dendritic cells from HAM/TSP patients show lower IFNα responsiveness

Several studies have reported alterations in cell frequencies between asymptomatic carriers and HAM/TSP subjects and increase of pro-inflammatory cytokines^10,12^. However, none of these studies have addressed alteration in cell responsiveness as a potential signature of the disease progression. Aiming at investigating this question we stimulated the blood collected from HTLV-1-asymtomatic subjects with the TLR7/8 agonist R848, also known as Resiquimod and performed immunophenotypical analysis of the innate cell populations by flow cytometry. Dendritic cells were gated from HLA-DR^+^ subset and classified as cDC1, cDC2 and pDC based on differential expression of CD11c, BDCA2/3 and BDCA1 (see Supplementary Figure 1). We found no differences in cDC1 and pDC cell frequencies between the clinical groups and controls. However, we found higher frequencies of cDC2 subset in HAM/TSP patients compared to AC (Figure 2A). Next, we investigated the cellular heterogeneity of innate cells using an unbiased high-dimensional analysis, with the aim to reveal subtle differences in multiple cell populations that may have been missed by the use of biaxial gating. tSNE analysis was applied to similar number of 15000 cells from all individuals in HD, AC and HAM/TSP groups (Figure 2B). This approach generates a two-dimensional map where similar cells are placed at adjacent points, while cells with different characteristics are separated in space. tSNE analysis showed differences in cDC1 population between clinical groups (Figure 2B). In fact, a small population of cDC1 was identified by tSNE analysis in healthy donors and AC at steady state. However, this population was practically disappeared in HAM/TSP patients. Interestingly, only HTLV-1-asymptomatic subjects (AC) maintained this population after TLR7 stimulation with R848. Intracellular readout of for IFNα, IL-12, Mip-1α and TNFα was performed in the different DC subsets. The frequency of cells producing these cytokines in each DC subset was represented in Figure 2C for both untreated and whole blood treated with R848 in HTLV-1-asymptomatic carriers (AC) and HAM/TSP patients. Resiquimod or R848 is an agonist of the TLR7/8 signaling pathway inducing the activation of dendritic cells and monocytes. Overall, we found a correct activation of all DC subsets. However, we found an impaired responsiveness of HAM/TSP blood samples (Figure 2C). Notably, IFNα production was significantly decreased in both cDC1 and cDC2. cDC1 had also lower production of TNFα and Mip-1α production was impaired in cDC2. In contrast, higher responsiveness to produce IL-12 was observed in pDC subset f HAM/TSP subjects.

Boolean analysis of the cytokine co-production showed an impaired responsiveness of the TNFα production by cDC1 and cDC2, as previously reported, as well as co-production of IFNα-TNFα by pDC (Figure 2D). In contrast, co-production of IL12-TNFα by cDC2 was significantly decreased in HAM/TSP compare to asymptomatic controls. Taken altogether, these data suggest that HAM/TSP patients present an impaired responsiveness of IFNα producing subsets, important for the antiviral response, while responsiveness of IL-12 subsets is increased.

**Figure 2.**
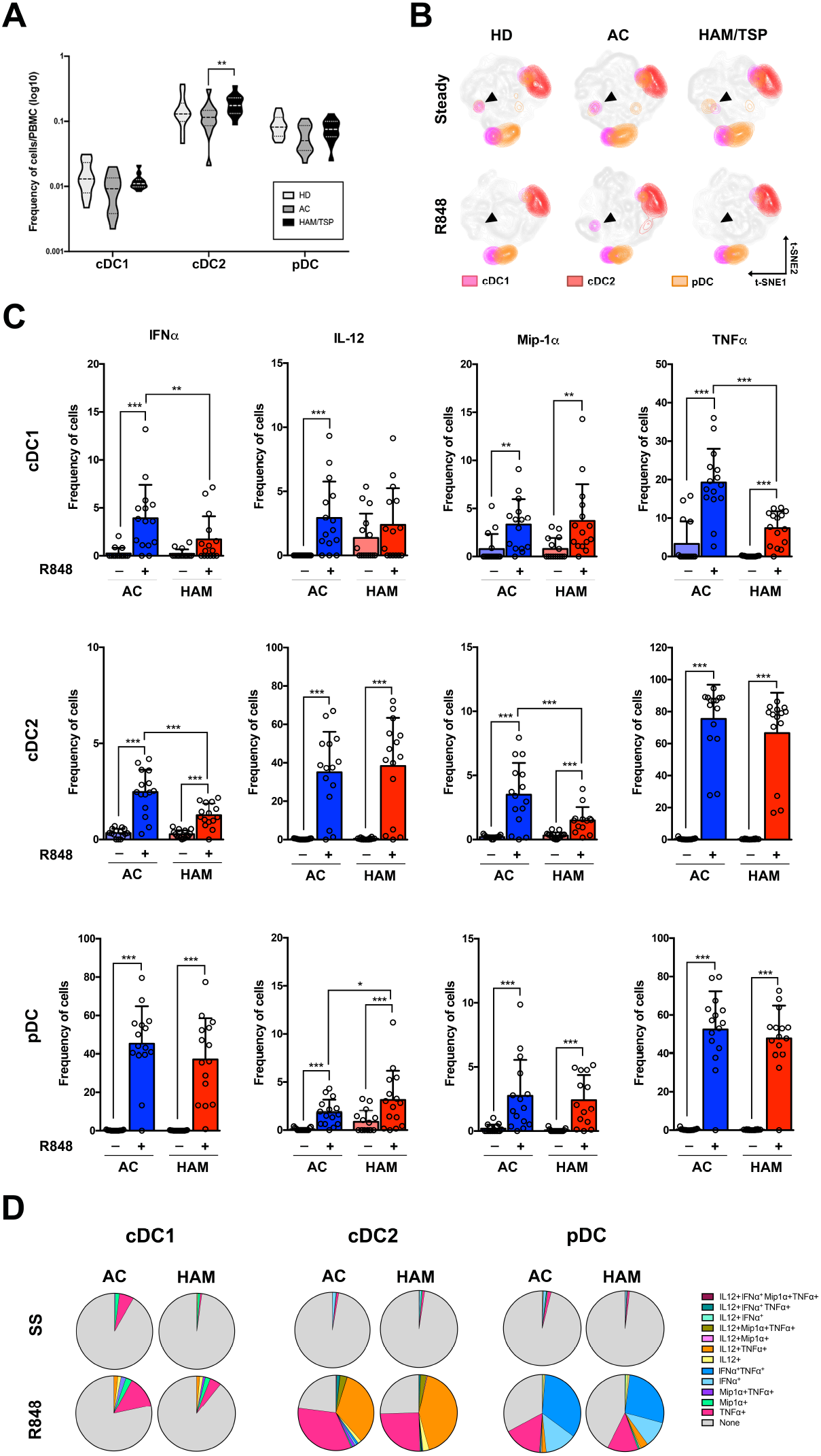
**(A)** Frequency of innate subsets in the clinical groups evaluated in dendritic cell subsets (cDC1, cDC2 and pDC). **(B)** tSNE analysis of DC subset frequencies in whole blood samples at steady state and after TLR7-stimulation with R848. **(C)** Frequency of the cells producing IFNα, IL-12, Mip-1α and TNFα at steady state and after R848 treatment in the different DC subsets. HD: healthy donor; AC: asymptomatic carriers; and HAM/TSP: HTLV-1 associated myelopathy/Tropical spastic paraparesis. * p-value ≤ 0.05; ** p-value ≤ 0.01; *** p-value ≤ 0.001. **(D)** Pie-chart of the Boolean analysis for the cytokine production in DC subsets at steady state (SS) and after TLR7 stimulation (R848).

### Monocytes from HAM/TSP patients show higher IL-12 and Mip-1α responsiveness

Monocytes were gated from HLADR^+^CD11c^+^ subset excluding BDCA1 expression and subsequently divided in three subpopulations based on the expression of CD16 and CD14 markers as: classical monocytes (cMono; CD14^+^CD16^-^), intermediate monocytes (intMono; CD14^+^CD16^+^) and non-classical monocytes (ncMono; CD14^-^CD16^+^) (Supplementary Figure 1 and Figure 3A). Consistent with a previous report^12^, a higher frequency of intermediate monocytes was detected in HAM/TSP patients compared to asymptomatic carriers (Figure 3B). tSNE distribution revealed subtle differences were found in the distribution of classical and intermediate monocytes between the three clinical groups (Figure 3C). Two clusters of intermediate monocytes were identified based on t-SNE pots, unequally distributed between AC and HAM/TSP especially after TLR7 stimulation.

**Figure 3.**
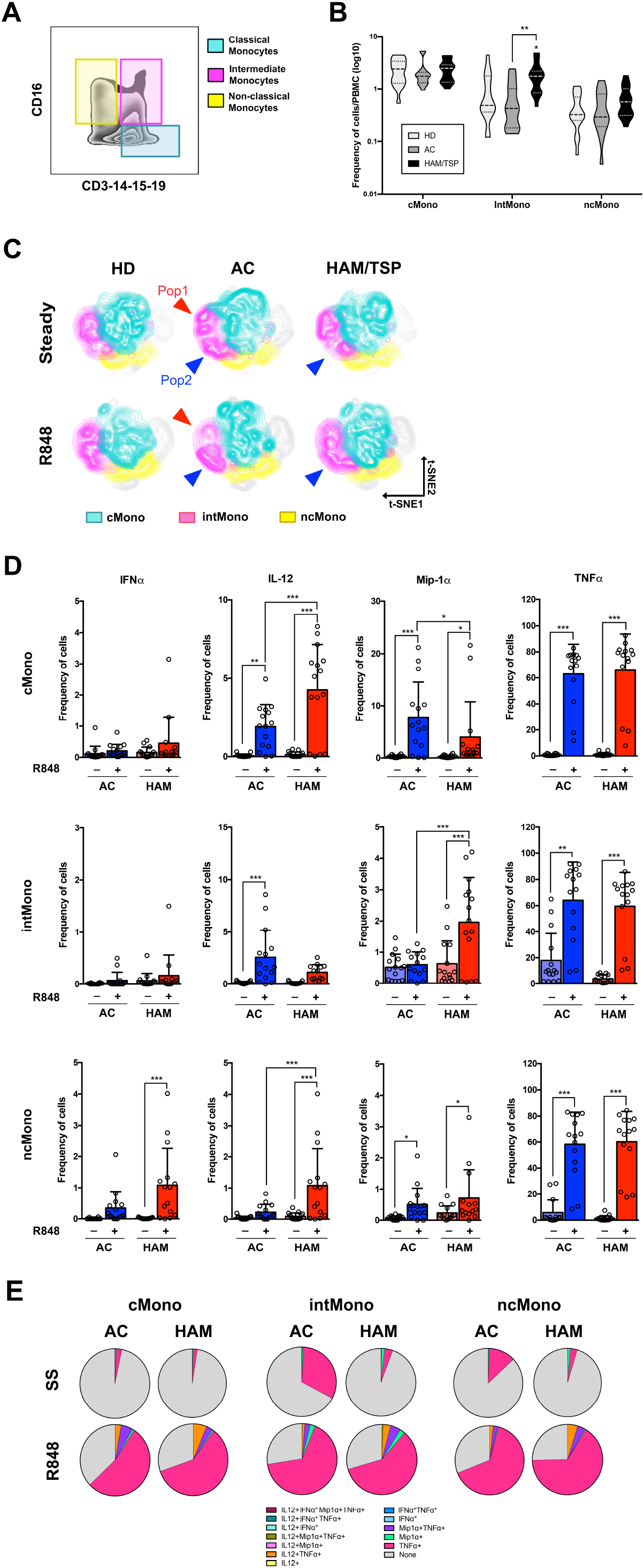
**(A)** Gating strategy for identification of the three monocyte subpopulation using CD16 and CD3-14-15-19 antibodies. **(B)** Frequency of innate subsets in the clinical groups evaluated in monocytes subsets (classical monocytes, intermediate monocytes and non-classical monocytes). **(C)** tSNE analysis of monocyte subset frequencies in whole blood samples at steady state and after TLR7-stimulation with R848. **(D)** Frequency of the cells producing IFNα, IL-12, Mip-1α and TNFα at steady state and after R848 treatment in the different monocyte subsets. HD: healthy donor; AC: asymptomatic carriers; and HAM/TSP: HTLV-1 associated myelopathy/Tropical spastic paraparesis. * p-value ≤ 0.05; ** p-value ≤ 0.01; *** p-value ≤ 0.001. **(D)** Pie-chart of the Boolean analysis for the cytokine production in monocyte subsets at steady state (SS) and after TLR7 stimulation (R848).

In contrast to what observed in dendritic cells, we found overall greater responsiveness of HAM/TSP monocytes to TLR7/8 stimulation (Figure 3D). Frequencies of IL-12-producing classical and non-classical monocytes and also Mip-1α intermediate monocytes were significantly higher in HAM/TSP subjects than in asymptomatic carriers. In contrast, the frequency of Mip-1α classical monocytes was lower in HAM/TSP compared to AC. The spontaneous production of TNFα was significantly decreased in HAM/TSP patients compared to AC (Figure 3E). After R848 stimulation the co-production of IL-12-TNFα was higher in HAM/TSP compared to AC.

Increased released of Mip-1α and IL-12 together with increased number of intermediate monocytes could mean a significant increase in the overall production of both cytokines. Increased IL-12 was linked with autoimmunity and can induce the expression of IFNγ and TNFα by NK and T-cells^22^.

### Higher IFNγ production found in NK cells from HAM/TSP

Natural Killer cells are though to make the bond between the innate and the adaptive immune response^23^, especially in autoimmune disease^24^. Furthermore, NK cell population of HTLV-1 infected subjects showed spontaneous proliferation capacity^25^. In this line, we aimed at investigating potential frequency alterations in our cohort of HTLV-1 infected donors. NK cells subsets were gated from HLA-DR^-^ subset (see Supplementary Figure 1) and divided into CD56^dim^CD16^+^ and CD56^high^CD16^-^ (Figure 4A). We observed a decreased in asymptomatic carriers of both NK subsets cell frequency compared to healthy donors and HAM/TSP patients presented lower cell frequencies of CD56^high^CD16^-^ NK cells compared to asymptomatic carriers (Figure 4B). tSNE distribution evinced an extra subpopulation of CD56^dim^CD16^+^ NK cells in asymptomatic carriers (Pop1) (Figure 4C), presenting high production of TNFα (Figure 4D). A small subpopulation of CD56^dim^CD16^-^ NK cells disappeared in asymptomatic carriers after TLR7 stimulation.

**Figure 4.**
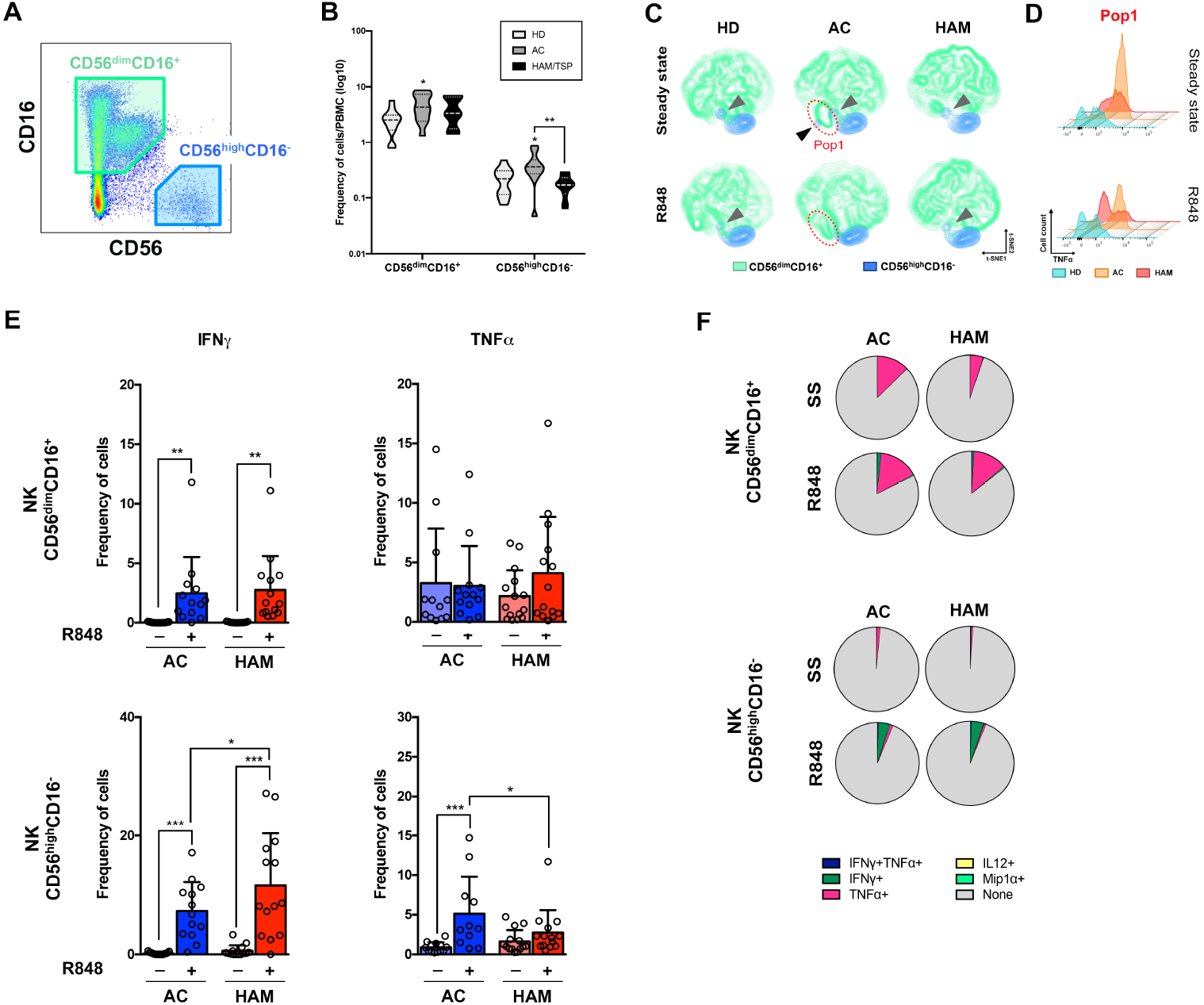
**(A)** Gating strategy followed to isolate NK cell populations according to CD16 and CD56 expression. **(B)** Violing-plot representation of the NK subpopulations cell frequency in the 3 clinical groups. **(C)** t-SNE clustering of the NK cells reveals different subpopulations and **(D)** the histogram represents TNFα expression of the identified population (Pop1). **(E)** Bar-plot representing the frequency of cytokine producing cells by the gated NK subpopulations at steady state and upon R848 stimulation. **(F)** Boolean analysis of the multiple-cytokine production by the NK subtypes at steady state (NS) and after TLR7/8 stimulation (R848).

The interaction with macrophages, lymphocytes and dendritic cells through response and secretion a variety of cytokines mediates the susceptibility and/or protective role of NK from disease progression^24^. In view of this fact, we wondered if the observed alteration in dendritic cells and monocytes immune response could trigger an NK dysfunctional response. While we found no significant differences in CD56^dim^CD16^+^ NK cells subset responsiveness, we observed that CD56^dim^CD16^-^ NK cells from HAM/TSP patients presented higher IFNγ production after TLR7 stimulation (Figure 4E). In contrast, the production of TNFα was significantly decreased in HAM/TSP compared to asymptomatic carriers. However the production of TNFα, especially by CD56^dim^CD16^+^ NK cells was already high in HTLV-1-asymtomatic carriers (Figure 4F). The low TNFα spontaneous production in HAM/TSP patients could be as a result of corticosteroid treatment.

## Discussion

HAM/TSP only occurs in 1-5% of HTLV-1-infected subjects but it represents a devastating neurodegenerative disease. To date, predictive markers of the evolution towards HAM/TSP pathology in HTLV-1 asymptomatic carriers is still a challenging task. We provided here that neither an elevated PVL –an average hallmark of HAM/TSP patients compared to asymptomatic carriers^21^– nor immune cell frequency alterations are sufficient to anticipate the disease progression. However, since innate compartment is also targeted by HTLV-1 infection –potentially contributing to functional deregulations– the unexplored innate response alterations could shed some light on the HAM/TSP pathology.

Due the rare frequency of HAM/TSP disease and the logistical difficulties of collecting and analyzing freshly blood samples, most of the up-to-date performed studies departed from cryopreserved samples, potentially affecting blood cellular components. However, even the detection of cell surface markers look unaffected^13^, we have observed an impaired immune response notably by dendritic cells and monocytes upon TLR7/8 signaling pathway stimulation. In addition, working with scarce frequencies such as those of dendritic cells could significantly affect cell counts if cell survival would be affected upon cryopreservation. In regard of these facts, our work has been performed on freshly collected blood samples of a cohort of 45 Brazilian volunteers. Aiming at identifying potential immune biomarkers that could anticipate symptomatic manifestations, we performed an immunoprofiling of innate cells functionality both at steady state and after stimulation of fresh blood samples collected from a cohort of 30 HTLV-1-infected subjects.

Our results suggest that HAM/TSP patients presented a diminished antiviral response by dendritic cells as IFNα levels produced after TLR7/8 stimulation were lower compared to asymptomatic carriers. This decreased antiviral response could be linked with increased PVL accumulating across time. The consistent decreased TNFα production found at steady state in HAM/TSP samples might be related to masked inflammatory profile as a result of corticosteroid treatment. However, by addressing responsiveness through TLR7/8 stimulation we observed greater responsiveness of HAM/TSP monocytes to produce IL-12 and Mip-1α. Mip-1α is an inflammatory chemokine, also named CCL3, has been associated with multiple sclerosis^26^, an autoimmune and inflammatory disease that affects brain and spinal cord functions, through demyelination of nerves, causing irreversible damages of the CNS. Mip-1α stimulates T-cell chemotaxis from periphery to inflamed tissues and regulates the transendothelial migration of monocytes, dendritic cells and NK cells^27^. Thus, the increased responsiveness of monocytes towards the production of Mip-1α in HAM/TSP individuals could be linked with neuronal inflammation through a favored invasion of immune cells in the CNS. Interestingly, potent antagonists of CCL3 receptors, CCR1 and CCR5, have been developed^28^ and their efficacy evaluated in clinical trials against multiple sclerosis among other inflammatory diseases. Thus use of such CCR1 antagonists might also be considered in the treatment of HTLV-1 infected individuals at risk of HAM/TSP.

On the other hand, higher responsiveness detected in some DC subsets and monocytes to produce IL-12 in HAM/TSP patients could contribute to the maintenance of an adaptive inflammatory response, despite anti-inflammatory treatment, as IL-12 strongly synergizes with other stimuli to induce a maximal production of IFNγ^29,30^ in T-cells and enhance NK cells cytotoxic activity^31^. In fact, we detected greater IFNγ production by NK cells in HAM/TSP patients after stimulation. Higher levels of spontaneous IFNγ produced by NK cells were detected in HAM/TSP patients correlating with a continuous activation state (Queiroz 2019). However, the pro-inflammatory profile seems to be extensive to all HTLV-1-infected individuals, deregulations in innate subsets observed here could be indicative of evolution towards HAM/TSP.

## Acknowledgements

BR would like to acknowledge Dr. Yamila Rocca (CIRI, T Walzer team). Authors specifically thank Sebastien Dessurgey and Thibault Andrieu (SFR biosciences, cytometry platform) for their technical advices and Dr. Chloé Journo for her critical reading, helpful discussions and her help in the statistics. Authors are also grateful to Dr. David Karlin for his advices in writing and to Dr Patrick Lecine for critical reading.

## Authorship Contributions

NF performed the experiments with the help of NO. BR analyzed the results, interpreted the data and wrote the first drafts of the manuscript. OA performed biostatistical analysis. JC and ACPO created the cohort, followed the patients in clinic and recruited the individuals analyzed in this study. RM supervised the study. HD conceptualized, supervised and directed the study. All authors discussed the results and commented on the manuscript.

## Supplementary Tables and Figures

**Supplementary Table 1.** Clinical information of the cohort. Table containing information about the HTLV-1-infected subjects enroller in the study. Clinical status, sex, age, PVL, motors score, treatment and breastfeeding information is detailed for each subject.

| Number | Clinical status | Age | Sex | PVL | Osame Score | IPEC Score | Treatment | Breastfeeding | Familiar situation |
| --- | --- | --- | --- | --- | --- | --- | --- | --- | --- |
| 1 | HAM/TSP | 52 | M | 205.81 | 6 | 14 | Yes | 12 months | Partner HTLV1+ |
| 2 | HAM/TSP | 52 | F | 477.26 | 4 | 12 | Yes | n/a | Divorced |
| 3 | HAM/TSP | 55 | M | 408.99 | 7 | 36 | Yes | Yes | Partner HTLV1+ |
| 4 | HAM/TSP | 55 | M | 88.98 | 8 | 22 | Yes | n/a | Married |
| 5 | HAM/TSP | 66 | F | 6084.07 | 9 | 23 | Yes | n/a | Widow |
| 6 | AC | 52 | F | 585.67 | n/a | n/a | No | 12 months | Partner HTLV1 Neg |
| 7 | AC | 35 | F | 619.39 | n/a | n/a | No | 6 months | Partner HTLV1 Neg |
| 8 | AC | 70 | F | 4.77 | n/a | n/a | No | Yes | Single |
| 9 | AC | 58 | F | 546.49 | n/a | n/a | No | n/a | Widow |
| 10 | AC | 46 | F | 480.33 | n/a | n/a | No | Yes | Divorced |
| 11 | AC | 41 | F | 1.39 | n/a | n/a | No | n/a | Single |
| 12 | AC | 18 | F | 20.14 | n/a | n/a | No | 24 months | Single |
| 13 | AC | 52 | M | 1.00 | n/a | n/a | No | n/a | Single |
| 15 | HAM/TSP | 45 | F | 927.13 | 7 | 20 | Yes | Yes | Partner HTLV1 Neg |
| 16 | HAM/TSP | 58 | F | 88.97 | 5 | 20 | Yes | n/a | Single |
| 17 | AC | 42 | F | 10.3 | n/a | n/a | No | 36 months | Partner HTLV1 Neg |
| 18 | AC | 46 | F | 2186.64 | n/a | n/a | No | n/a | n/a |
| 19 | HAM/TSP | 46 | F | 274.94 | 5 | 15 | Yes | 24 months | Partner HTLV1 Neg |
| 21 | AC | 44 | F | 1.00 | n/a | n/a | No | Yes | Widow |
| 28 | AC | 36 | M | 112.04 | n/a | n/a | No | n/a | Divorced |
| 29 | HAM/TSP | 57 | F | 1.88 | 8 | 18 | Yes | n/a | Single |
| 30 | HAM/TSP | 60 | F | 434.89 | 5 | 14 | Yes | Yes | Divorced |
| 31 | HAM/TSP | 54 | M | 565.15 | 7 | 23 | Yes | n/a | Partner HTLV1 Neg |
| 32 | HAM/TSP | 52 | F | 1057.04 | 5 | 14 | Yes | n/a | Partner HTLV1 Neg |
| 33 | HAM/TSP | 59 | M | 3085.02 | 6 | 15 | Yes | 96 months | Partner HTLV1 Neg |
| 35 | AC | 53 | F | 1.0 | n/a | n/a | No | n/a | Partner HTLV1 Neg |
| 43 | HAM/TSP | 63 | F | 118.53 | 5 | n/a | Yes | n/a | Divorced |
| 44 | HAM/TSP | 34 | F | 679.99 | 5 | 15 | Yes | Yes | Single |
| 48 | AC | 46 | F | 42.41 | n/a | n/a | No | 48 months | Partner HTLV1+ |
| 49 | AC | 65 | F | 319.77 | n/a | n/a | No | Yes | Partner HTLV1 Neg |

**Supplementary Table 2.** List of antibodies. Recapitulative list of the **(A)** membrane markers antibodies and **(B)** intracellular markers antibodies used for the analysis of the innate immune response by flow cytometry.

**A**
| Antibody | Clone | Fluorochrome | Supplier |
| --- | --- | --- | --- |
| CD3 | UCHT1 | BV510 | BD Bioscience |
| CD14 | M5E2 |  |  |
| CD15 | HI98 |  |  |
| CD19 | HIB19 |  |  |
| HLA-DR | G46-6 | APC-H7 |  |
| CD11c | Bu15 | PerCP-Cy5.5 | Biologend |
| BDCA-2 | AC144 | BV421 | Miltenyi |
| BDCA-3 | AD5-14H12 |  |  |
| BDCA-1 | L161 | PE-Cy7 | Biologend |
| CD56 | HCD56 | BV605 | BD Bioscience |

**B**
| Antibody | Clone | Fluorochrome | Supplier |
| --- | --- | --- | --- |
| INFα | LT27:295 | FITC | Miltenyi |
| INFγ | 4S.B3 | BV786 | Biologend |
| IL-12p40/70 | C11.5 | PE | BD Bioscience |
| Mip-1α | 11A3 | APC |  |
| TNFα | Mab11 | AF700 |  |

**Supplementary Figure 1.**
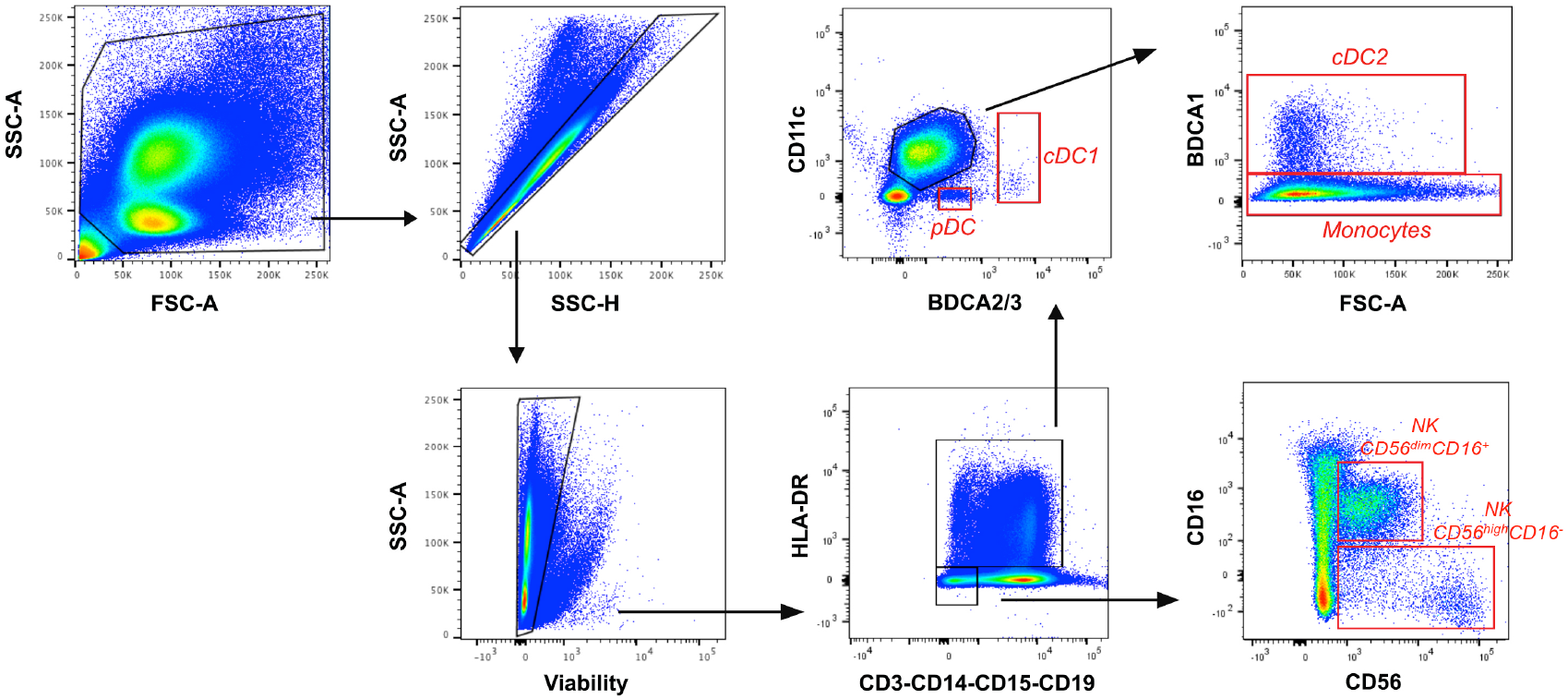
Hierarchical gating strategy of the different immune cell subpopulations. Flow cytometry collected datasets were analyzed with FlowJo software. A total of 2×10^6^ cells were registered and selected by cell size and granularity. After selection of single cells, viable cells were gated and innate immune cell populations were identified as indicated.

**Supplementary Figure 2.**
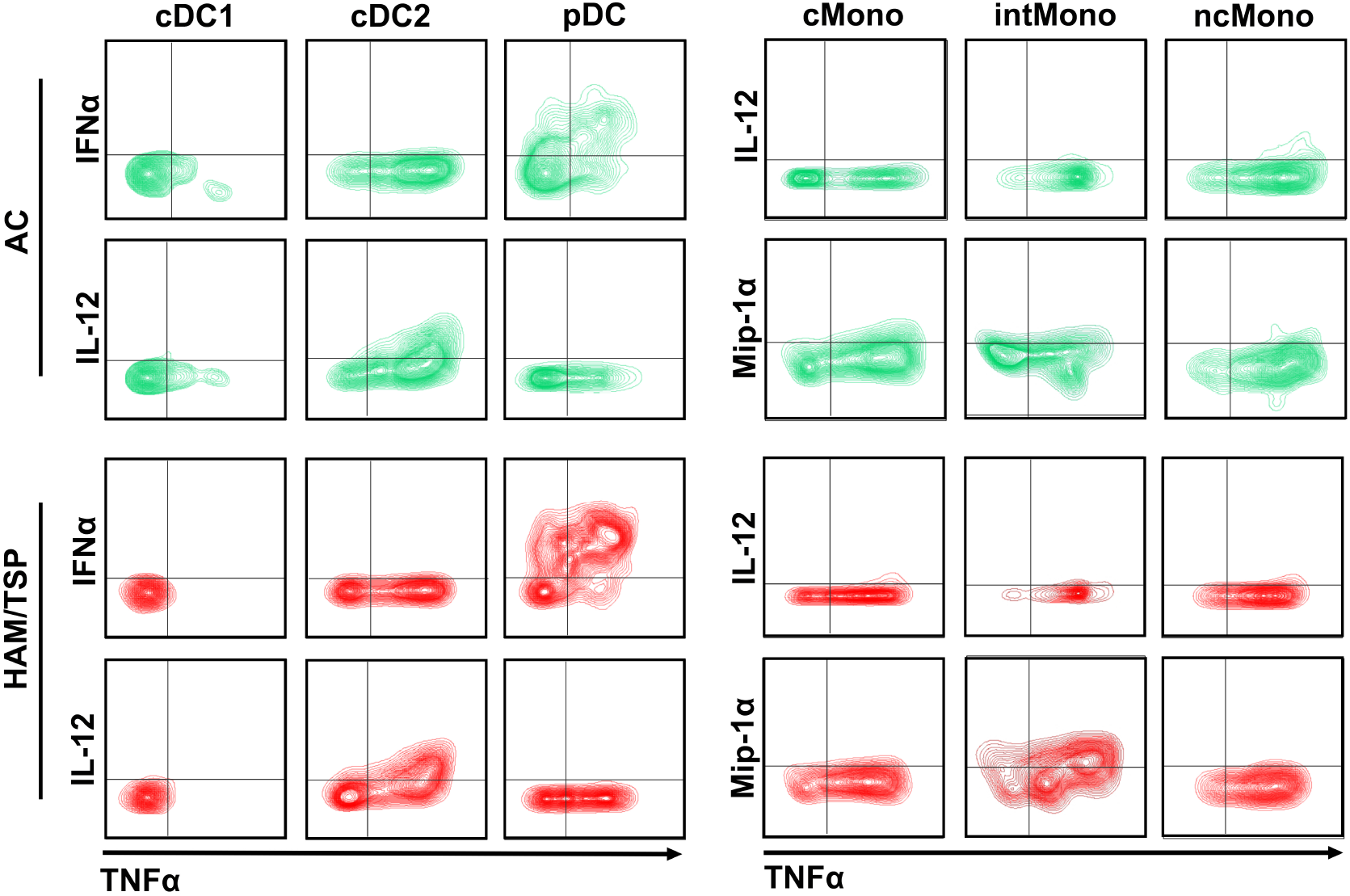
Gating strategy for cytokines. Example of the sating strategy for cytokine determination in AC and HAM/TSP group for IFNα, IL-12 Mip-1α and TNFα in the different cell subsets.

